# Validation and identification of new QTLs for plant and fruit developmental and composition traits in eggplant under low N conditions

**DOI:** 10.1101/2024.05.06.592694

**Authors:** Gloria Villanueva, Santiago Vilanova, Mariola Plazas, Pietro Gramazio, Jaime Prohens

## Abstract

Enhancing plant adaptation to low input conditions is a fundamental goal for implementing sustainable agriculture. In the present study, two eggplant (*Solanum melongena*) accessions (MEL1 and MEL5), two introgression lines (ILs) derived from eggplant wild relatives *S. dasyphyllum* (IL-M1-D1) and *S. insanum* (IL-M5-I9), and a heterozygous version of this last IL (IL^Het^-M5-I9), along with hybrids among them were evaluated under low N (LN) conditions. IL-M1-D1 carries an introgressed fragment of 4.9 Mb in homozygosis from *S. dasyphyllum* on chromosome 2, while IL-M5-I9 and IL^Het^-M5-I9 carry an introgression of 21.5 Mb on chromosome 9 in homozygosis and heterozygosis, respectively, from *S. insanum*. Multiple quantitative trait loci (QTLs) for several traits of interest were associated with both introgressions under LN conditions in a previous study with segregating advanced backcrosses. Here we evaluated the performance of these materials for 22 agronomic and developmental traits under low N fertilization (LN) conditions. Hybrids with the ILs enabled the study of genetic background effects on QTLs expression. The materials evaluated showed a significant phenotypic variation, particularly within hybrids segregating for the introgression from *S. insanum* in chromosome 9. Statistical analysis revealed no significant differences among hybrids carrying or not the introgression on chromosome 2 of *S. dasyphyllum*, and only slight differences were observed between the IL-M1-D1 and its recurrent parent *S. melongena* MEL1, suggesting a limited impact of this introgression on chromosome 2 on the phenotype variation. However, the differences observed between IL-M5-I9 and its recurrent parent *S. melongena* MEL5, together with the association between genotypic and phenotypic variation in hybrids segregating for this introgression, allowed the identification of 13 QTLs on chromosome 9. These results successfully validated the previously identified QTLs for flavonol content in leaves, nitrogen balanced index, fruit mean weight, and nitrogen content in leaves and, also revealed nine new QTLs associated with the introgressed genomic region in chromosome 9. This study emphasizes the influence of environmental conditions, genotypes, and genetic backgrounds on the phenotypic expression of eggplant QTLs introgressed from wild relatives and highlights the importance of QTL validation. These findings contribute valuable insights for developing new eggplant cultivars for a more sustainable agriculture, particularly with adaptation to LN conditions.

## 1. Introduction

In recent years, the impact of climate change has led to increased abiotic stress conditions such as drought, salinity, extreme temperatures, and nutrient deficiencies. These factors are limiting the use of arable land globally and negatively affect crop productivity (Pascual et al., 2022). Consequently, finding new strategies to enhance sustainable crop production and resilience has become a main objective in plant breeding (Bailey-Serres et al., 2019; Zhang et al., 2022). While nitrogen (N) fertilization is commonly used to improve crop yields, its overuse and inefficiency can increase production costs and cause considerable environmental damage (Stevens, 2019).

Eggplant (*Solanum melongena* L.), also known as brinjal or aubergine, ranks as the second most important vegetable crop in global production after tomato within the Solanaceae family (FAOSTAT, 2023). Cultivated eggplant exhibits a large diversity of morpho-agronomic traits, which is highly valuable for breeding purposes (Taher et al., 2017). However, there is a limitation in its diversity within the cultivated genepool, especially in traits for adaptation to climate change (Plazas et al., 2019). Crop wild relatives (CWRs) of eggplant, comprising over 500 species of *Solanum* subgenus *Leptostemonum* (Knapp et al., 2019) across its primary, secondary and tertiary genepools, represent a valuable source of plant genetic resources adapted to diverse stressful environments (Dempewolf et al., 2017; Prohens et al., 2017). This extensive genetic diversity in eggplant wild relatives is fundamental in creating new genetic resources and developing improved varieties for a more sustainable agriculture in a climate change scenario (Gramazio et al., 2023; Plazas et al., 2020; Toppino et al., 2022). Belonging to the primary genepool (GP1), *S. insanum* L. is considered the wild progenitor of the common eggplant (*S. melongena*) (Ranil et al., 2017). *Solanum dasyphyllum* Schumach. and Thonn. is considered the wild ancestor of the African gboma eggplant (*S. macrocarpon* L.) and it belongs to the secondary genepool (GP2) of eggplant (Taher et al., 2017; Vorontsova and Knapp, 2016). Both wild species display adaptability to various environmental conditions and have been reported to display tolerance to abiotic stresses such as drought and salinity (Kouassi et al., 2020; Ortega-Albero et al., 2023; Villanueva et al., 2023b).

Recent advances in eggplant genomics have been marked by a notable increase in the development of high-quality genome assemblies (Barchi et al., 2021, 2019b; Li et al., 2021; Wei et al., 2020). Additionally, the development of high-throughput genotyping platforms (Barchi et al., 2019a) has enabled extensive genotyping of both eggplant and its wild relatives. This has facilitated the development of interspecific hybrids, advanced backcrosses (ABs), introgression lines (ILs), and MAGIC populations in eggplant breeding (García-Fortea et al., 2019; Gramazio et al., 2017; Kouassi et al., 2016; Mangino et al., 2022; Plazas et al., 2016; Villanueva et al., 2021). In this way, the association of phenotypic and genotypic variation in eggplant has been crucial in identifying quantitative trait loci (QTLs) and candidate genes associated with a broad range of traits (Arrones et al., 2022; Mangino et al., 2021; Portis et al., 2014; Rosa-Martínez et al., 2023; Sulli et al., 2021; Toppino et al., 2020; Villanueva et al., 2023a).

In a previous study (Villanueva et al., 2023a), QTLs associated with flavonol content in leaves, nitrogen balanced index, fruit mean weight, and nitrogen content in leaves and stem (*fl-9, nb-9, fw-9, nl-9* and *ns-9*) were identified on a genomic region of chromosome 9 (7.4 Mb) in advanced backcrosses (ABs) derived from *S. insanum*. Additionally, another genomic region was detected on chromosome 2 (4.8 Mb) in ABs of *S. dasyphyllum*, carrying QTLs for chlorophyll leaf content, aerial plant biomass and yield (*ch-2, bi-2* and *yd-2*). This current study aimed to validate these QTLs previously identified in different genetic backgrounds, discover new QTLs of interest for eggplant breeding, and assess the potential for combining the QTLs present in both introgressions. For this purpose, we evaluated two *S. melongena* accessions (MEL1 and MEL5), two introgression lines (IL-M1-D2 and IL-M5-I9) derived from eggplant wild relatives *S. dasyphyllum* and *S. insanum* carrying alternative alleles for the QTLs, and segregating hybrids resulting from crosses of both introgressions. The results will allow us to assess the impact of genetic background and environmental factors on these traits and facilitate the development of N-efficient eggplant varieties.

## 2. Materials and methods

### 2.1 Plant material and growing conditions

For the study, two accessions of *S. melongena* (MEL1 and MEL5) were assessed, along with two introgression lines (IL-M1-D2 and IL-M5-I9) derived from MEL1 and MEL5, respectively. The IL-M1-D2 line, derived from *S. melongena* (MEL1), carries an introgressed fragment on chromosome 2 (6%; 4.9 MB) from the wild relative *S. dasyphyllum* (accession DAS1) with associated QTLs for chlorophyll content in leaf, aerial plant biomass, and yield (Villanueva et al., 2023a). On the other hand, the IL-M5-I9 was derived from *S. melongena* (MEL5) with an introgressed fragment on chromosome 9 (60%; 21.5 MB) from the wild relative *S. insanum* (accession INS1), containing QTLs for flavonol content in leaves, nitrogen balanced index, fruit mean weight, and nitrogen content in leaves and stem. Additionally, a version of IL-M5-I9 heterozygous for the introgression of the genomic region of chromosome 9 of *S. insanum* (IL^Het^-M5-I9) was used to obtain hybrids segregating for the introgression of *S. insanum* after crossing it with MEL1 and IL-M1-D2 (MEL1 X IL^Het^-M5-I9 and IL-M1-D2 X IL^Het^-M5-I9). The genetic background of these hybrids was characterized by heterozygosity for polymorphic loci between MEL1 and MEL5, as well as for segregating for the introgression of chromosome 9 of *S. insanum* (Figure 1). In total, the plant material consisted of 12 plants each of MEL1, MEL5, and IL-M1-D2, 11 plants of IL-M5-I9, 19 plants of segregating hybrids for the chromosome 9 introgression of *S. insanum* from MEL1 X IL^Het^-M5-I9, and 20 plants of segregating hybrids for the chromosome 9 introgression of *S. insanum* from IL-M1-D2 X IL^Het^-M5-I9.

**Figure 1.**
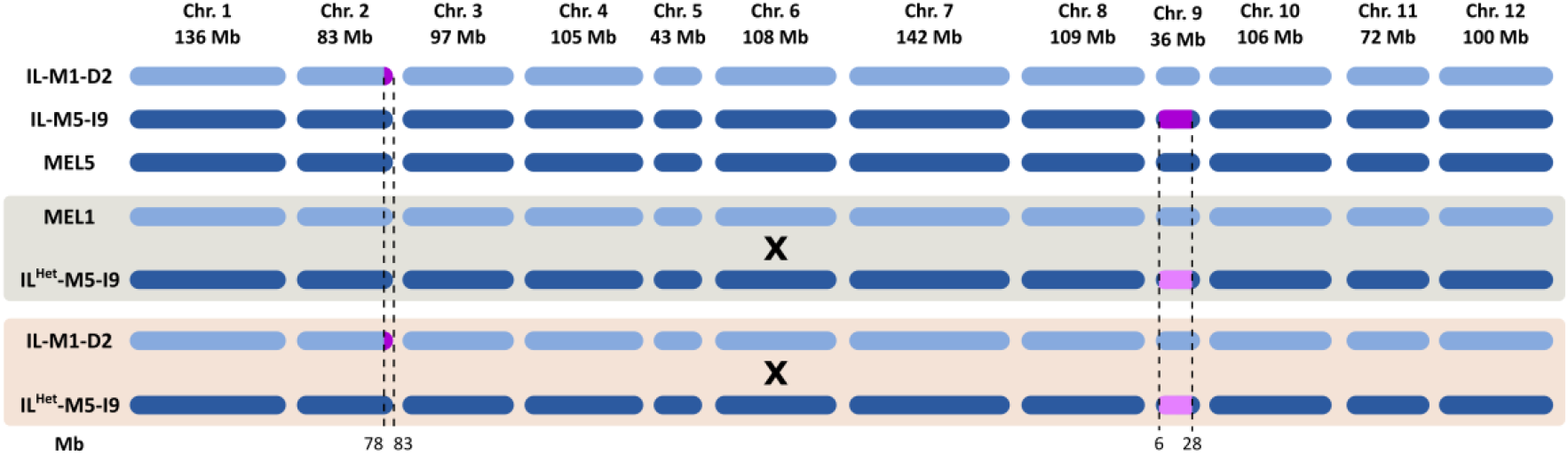
Graphical genotypes of introgression lines IL-M1-D2, IL-M5-I9, and MEL5, along with crosses MEL1 X IL^Het^-M5-I9 and IL-M1-D2 X IL^Het^-M5-I9. Columns represent the eggplant chromosomes and their respective length (Mb). Homozygous introgressions are colored in purple, heterozygous introgression in pink, and the genetic background of *S. melongena* MEL1 is colored in light blue, and *S. melongena* MEL5 in dark blue.

Plants were grown in an open field located in Universitat Politècnica de València (GPS coordinates: latitude, 39° 28’ 55” N; longitude, 0° 20’ 11” W; 7 m a.s.l.) during the summer season (July to October) of 2022. The experimental design involved the random distribution of plants in the field. Plants were grown in 17 L pots filled with coconut fiber. Irrigation and fertilization were applied using a drip irrigation system, under low nitrogen (LN) conditions.

The fertilization solutions were prepared based on the substrate composition and intake water analyses as detailed in Villanueva et al. (2023a). The LN fertilization solution was prepared by adding 1.5 mM H_3_PO_4_ (Antonio Tarazona SL., Valencia, Spain), 4.85 mM K_2_SO_4_ (Antonio Tarazona SL.), 0.58 mM MgSO_4_ (Antonio Tarazona SL.) plus 0.025 L/m^3^ of a microelements Welgro Hydroponic fertilizer (Química Massó S.A., Barcelona, Spain) containing boron (BO_3_^3-^; 0.65% p/v), copper (Cu-EDTA; 0.17% p/v), iron (Fe-DTPA; 3.00% p/v), manganese (Mn-EDTA, 1.87% p/v), molybdenum (MoO4^2^?; 0.15% p/v), and zinc (Zn-EDTA; 1.25% p/v). The pH of the irrigation solution was adjusted to 5.5–5.8 with 23% HCl (Julio Ortega SL., Valencia, Spain).

### 2.2 Genotyping

Genomic DNA extraction was performed for each individual plant using the SILEX DNA extraction method (Vilanova et al., 2020). High-throughput genotyping was conducted using the eggplant 5k probes single primer enrichment technology (SPET) platform (Barchi et al., 2019a). Single nucleotide polymorphisms (SNPs) were selected by filtering discriminant SNPs between the parental lines of introgression lines. Subsequently, a set of 824 SNPs were identified using TASSEL software (version 5.2.92; Bradbury et al., 2007).

### 2.3 Traits evaluation

A total of 22 plant, fruit and composition traits were evaluated. Chlorophyll, flavonol, anthocyanin contents, and nitrogen balanced index (NBI) were measured in leaves using a DUALEX® optical leaf clip meter (Force-A, Orsay, France). The dataset was obtained as the mean of 10 measurements of five leaves including upper and lower sides of each plant. At the end of the trial, aerial biomass was measured using a Sauter FK-250 dynamometer (Sauter, Balingen, Germany), and stem diameter was determined with a caliper at the base of the stem. Leaves and stems were separated and ground after drying at room temperature. The constant dry weight was measured after drying in an oven at 70ºC. Yield was determined by harvesting the total number of fruits from each plant. Nitrogen uptake efficiency (NUpE) was calculated by dividing the total N content in plant and fruit by the total N supplied with the irrigation solution per plant. Nitrogen utilization efficiency (NUtE) was calculated as total dry yield divided by the total N content in plant and fruit. Nitrogen use efficiency (NUE) was determined by multiplying NUpE with NutE (Anas et al., 2020; Han et al., 2015).

Fruit traits, including pedicel length, calyx length, fruit length, and width, were measured in fruits harvested at the commercially mature stage using a caliper. Data were calculated as the mean measurements obtained of at least three fruits per plant.

For the determination of nitrogen (N) and carbon (C) content, a minimum of five fruits per plant were harvested, frozen, and subsequently lyophilized. Analysis of N content in the dry powder from leaves, stem, and fruits was performed using the Dumas method with a TruSpec CN elemental analyzer (Leco, MI, USA). The measurement of carbon (C) content involved calculating carbon dioxide (CO2) using an infrared detector (Gazulla et al., 2012). The quantification process was performed with certified reference standards of different nitrogen and carbon concentrations with certified reference standards of different N and C concentrations.

### 2.4 Statistical analysis

For each trait, mean and range values were calculated for each parent, IL or segregating hybrid. To evaluate significant differences among recurrent parents, ILs and segregating hybrids, an analysis of variance (ANOVA) was performed. Significant differences were detected with the Student-Newman-Keuls multiple range test at a significance level of P < 0.05 using Statgraphics Centurion XIX software (Statgraphics Technologies, Inc. The Plains, VA, USA).

Principal component analysis (PCA) and partial least squares -discriminant analysis (PLS-DA) were performed for individual plant data using the package *ropls* (Thévenot et al., 2015) of the R software (R Core Team, 2021). The resulting PCA and PLS-DA plots were generated utilizing the *ggplot2* package (Wickham, 2016). PCA and PLS-DA are complementary multivariate techniques, in which PCA is a dimensionality reduction method based on maximizing the explanation of variance in the original variables, whereas PLS-DA is designed to reduce dimensionality by optimizing covariance between predictor matrix and a response matrix (Lee et al., 2018). In this approach, the PLS-DA was grounded in a predetermined group classification for each of the plant materials used, including MEL1, MEL5, IL-M1-D2, IL-M5-I9, MEL1 X IL^Het^-M5-I9, and IL-M1-D2 X IL^Het^-M5-I9.

### 2.5 Quantitative trait loci (QTL) detection

Identification of significant QTLs was performed using the single QTL model for genome-wide scanning with the R package *R/qtl* (Broman et al., 2003) of R statistical software (R Core Team, 2021). The threshold for significance in the logarithm of odds (LOD) score was established at a probability level of 0.01 for the significant QTLs. For each putative QTL detected, allelic effects were calculated by establishing differences between the means values of each genotype.

## 3. Results

### 3.1 Characterization of phenotypic traits

Significant differences among the different plant materials were found for all traits except for anthocyanin content in leaves (P-Anth), aerial plant biomass (P-Biomass), yield and NUE-related traits (NUE, NUpE and NUtE) (Table 1). No significant differences were observed between both hybrids derived from two crosses (MEL1 X IL^Het^-M5-I9 and IL-M1-D2 X IL^Het^-M5-I9) for any of the traits.

**Table 1.**
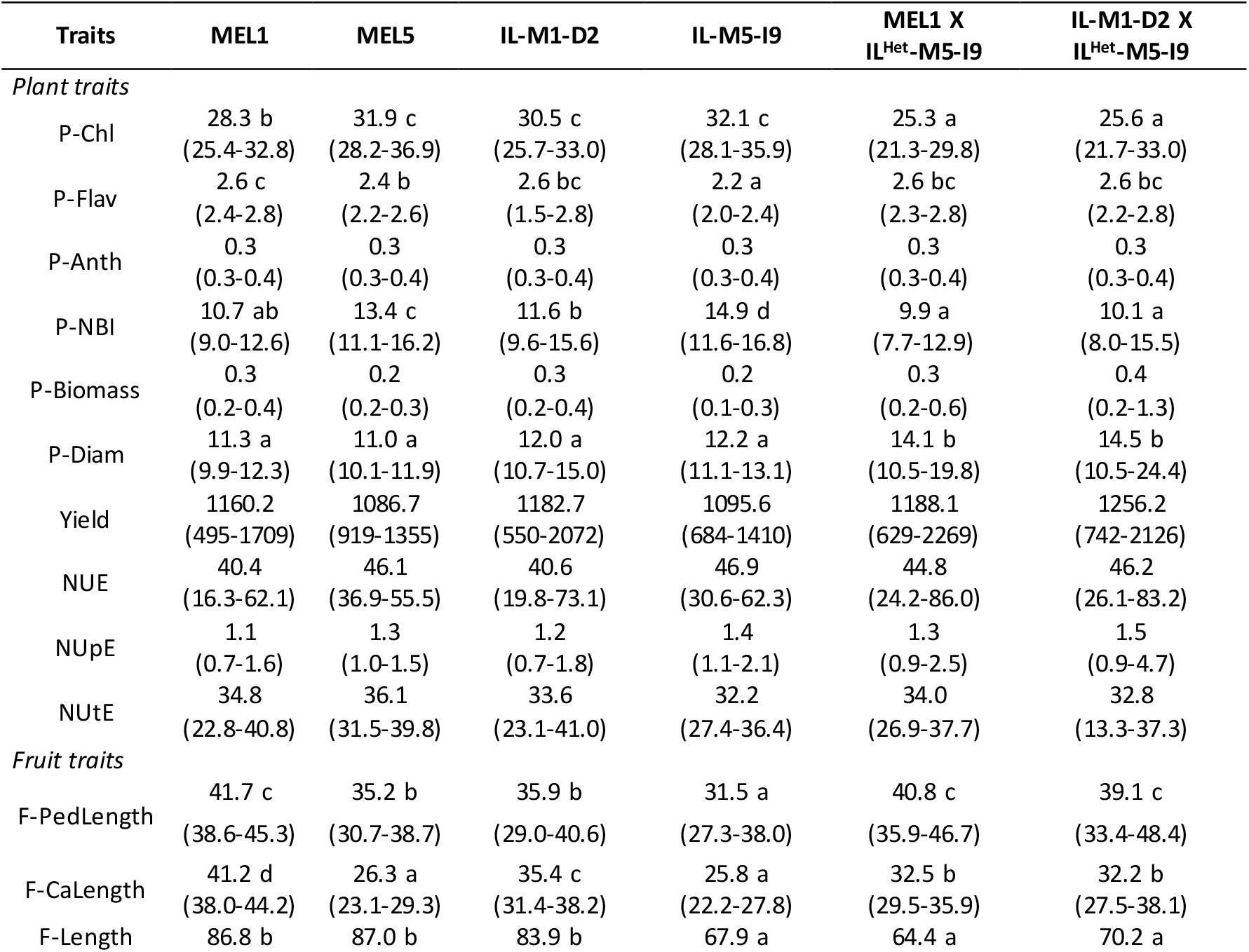
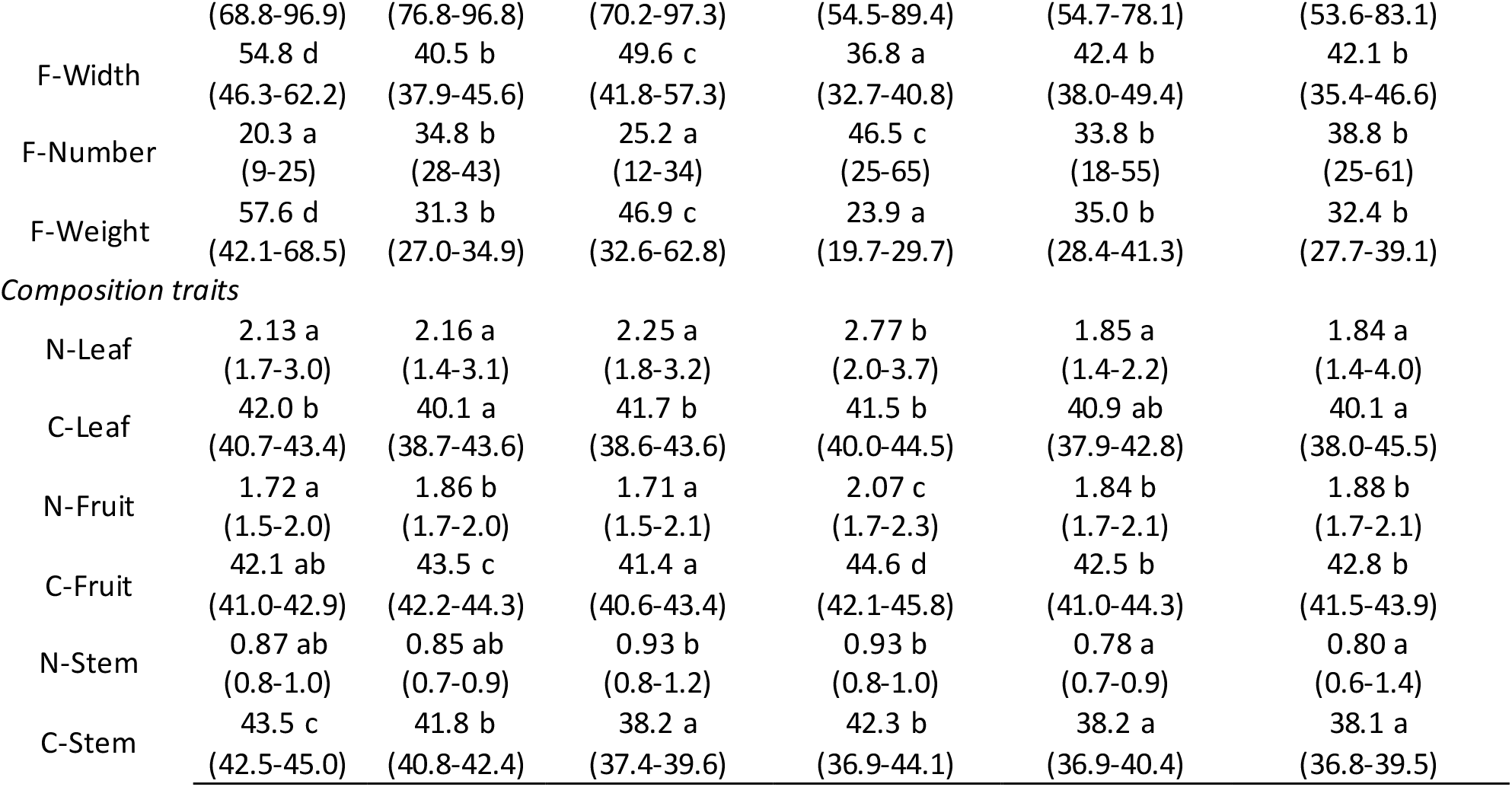
Mean values and range of plant, fruit and composition traits evaluated for *S. melongena* MEL1 (n=12), MEL5 (n=12), ILs of *S. dasyphyllum* IL-M1-D2 (n=12), ILs of *S. insanum* IL-M5-I9 (n=11), and segregating hybrids MEL1 X IL^Het^-M5-I9 (n=19) and IL-M1-D2 X IL^Het^-M5-I9 (n=20). For each trait, means with different letters are significant according to the Student–Newman– Keuls multiple range test (*P*<0.05).

For plant traits, significantly lower chlorophyll content in leaves (P-Chl) and nitrogen balanced index (P-NBI), along with a higher stem diameter (P-Diam), were observed in both segregating hybrids in contrast to other parents and ILs, indicating negative heterosis for these traits resulting from the cross between MEL1 and MEL5 genetic backgrounds. Additionally, the IL of *S. insanum* (IL-M5-I9) showed significantly lower flavonol content in leaves (P-Flav) than the rest of materials and the highest nitrogen balanced index (P-NBI) (Table 1).

In terms of fruit traits related to shape and size (F-PedLenght, F-CaLength, F-Length and F-Width), the parental line *S. melongena* MEL1 displayed significantly larger fruits, characterized by longer pedicels, calyx lengths, and greater fruit length and width, in comparison to the other accessions (Table 1). Conversely, the IL-M5-I9 introgression line exhibited significantly lower values for these four traits. Both segregating hybrids showed a significant higher fruit pedicel length (F-PedLength) together with *S. melongena* MEL1 and a significant lower fruit length (F-Length) along with IL-M5-I9 in contrast to other accessions. Furthermore, *S. melongena* MEL1 and IL-M1-D2 presented the significantly lowest number of fruits per plant (F-Number), while IL-MEL5-INS9 showed the highest number of fruits. Inversely, *S. melongena* MEL1 presented the highest mean fruit weight (F-Weight), and IL-M5-I9 displayed the lowest (Table 1).

Regarding composition traits, the accession IL-M5-I9 exhibited the significantly highest content of nitrogen (N) in leaves and fruits (N-Leaf and N-Fruit). On the contrary, *S. melongena* MEL1 and IL-M1-D2 showed the significantly lowest N content in fruits. Additionally, for carbon content in leaves (C-Leaf), both segregating hybrids, along with *S. melongena* MEL5, displayed significantly lower values, and for carbon content in stems (C-Stem), the segregating hybrids, along with IL-M1-D2, demonstrated significantly lower values compared to the other accessions (Table 1). The IL-M5-I9 displayed the significant highest carbon content in stem (C-Stem).

Overall, segregating hybrids derived from the IL-M1-D2 X IL^Het^-M5-I9 cross presented the widest ranges in most traits. Conversely, parental lines of *S. melongena* MEL1 and MEL5 displayed the narrowest ranges (Table 1).

### 3.2 Principal component and partial least squares analyses

A principal component analysis (PCA) was conducted for all evaluated traits, with a principal component (PC) 1 and PC2 accounting for 27.1% and 21.0% of the total variation, respectively (Figure 2A). The distribution pattern in the PCA revealed that the PC1 was positively correlated while the PC2 negatively correlated with nitrogen, and carbon content in fruit (N-Fruit and C-Fruit), nitrogen content in leaves (N-Leaf), chlorophyll content, and nitrogen balanced index in leaves (P-Chl and P-NBI), while. The first component was negatively correlated with leaf flavonol content (P-Flav), and traits related to fruit morphology and size (F-PedLength, F-CaLength, F-Length, F-Width and F-Weight). Regarding the individuals of the different plant materials, the IL-M5-I9 genotypes were predominantly located along the positive PC1 axis and negative PC2, while other genotypes generally displayed a wider distribution along the negative values of PC1 (Figure 2A).

**Figure 2.**
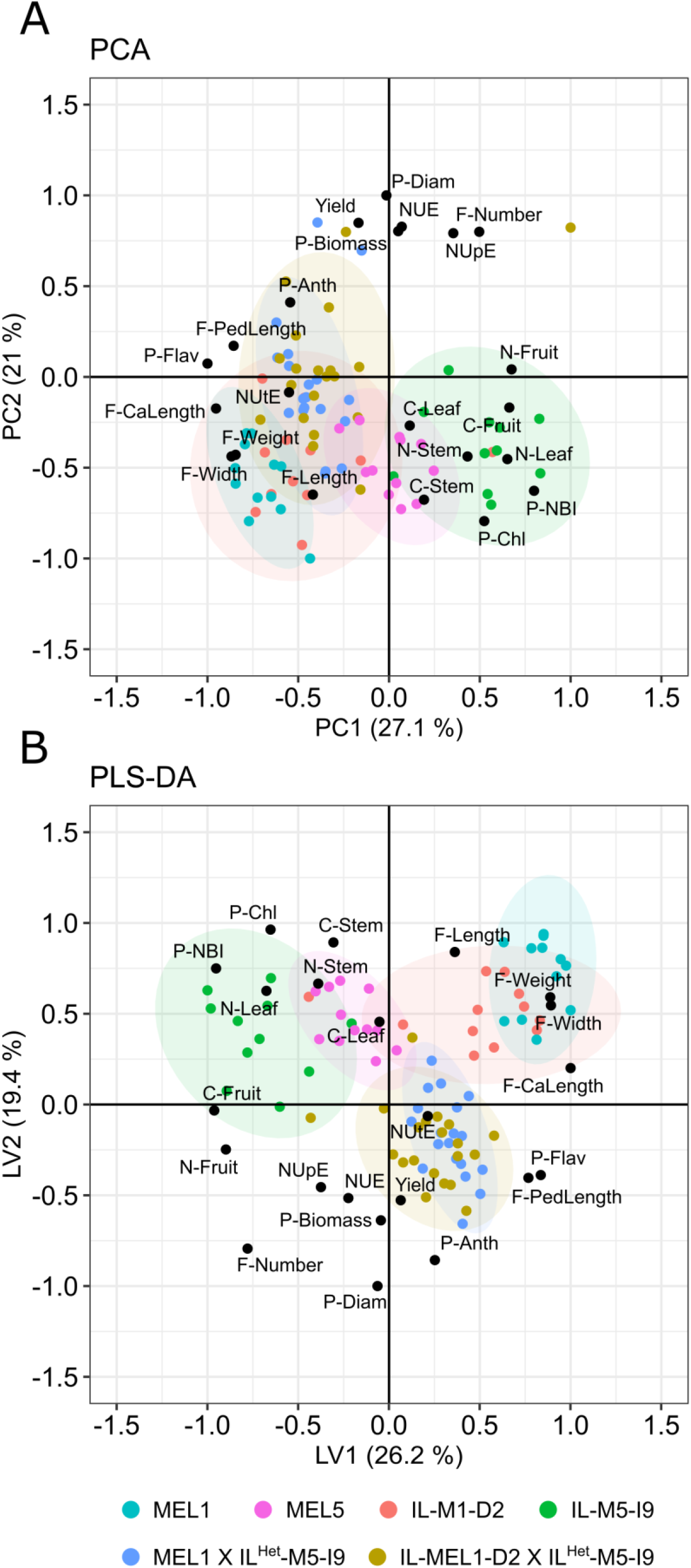
PCA (A) and PLS-DA (B) biplots based on the two principal components (PC1 and PC2) of the analysis performed for all traits.

The two first components (PCs) of the partial least squares analysis (PLS-DA) accounted for 26.2% and 19.4% of the total variation across all traits (Figure 2B). Traits related to fruit characteristics displayed positive values along the PC1 and PC2. The first component was negatively correlated with chlorophyll and nitrogen content in leaves (P-Chl and N-Leaf), and nitrogen balanced index (P-NBI). Additionally, flavonol and anthocyanin content in leaves (P-Flav and P-Anth), fruit pedicel length (F-PedLength), yield, and nitrogen utilization efficiency (NutE) had negative values for the PC1 axis and positive ones for the PC2. The distribution of the individuals in the PLS-DA plot revealed a distinct separation among the different materials. Specifically, the IL-M1-D2, along with the parental line *S. melongena* MEL1, mostly exhibited positive values along both PC1 and PC2. In contrast, IL-M5-I9 was predominantly positioned at negative values of PC1 and positive PC2 values, while *S. melongena* MEL5 was in its proximity. Hybrids resulting from two different crosses, MEL1 X IL^Het^-M5-I9 and IL-M1-D2 X IL^Het^-M5-I9, plotted very closely with generally positive values for the PC1 and negative ones for the PC2 (Figure 2B).

### 3.3 Detection and effects of putative QTLs

Genotypic and phenotypic association analyses facilitated the identification of 13 putative QTLs, all located at the same position (6.43 and 27.90 Mb) on chromosome 9 (Table 2). Regarding plant traits, QTLs associated with leaf chlorophyll content (*ch-9*) and balanced nitrogen index (*nb-9*) showed a significant positive effect of the homozygous allele of *S. insanum* (10.86% and 30.27%, respectively) with a contrasting negative effect for the heterozygote (-14.97% and -17.18%, respectively). Conversely, the QTL detected for leaf flavonol content (*fl-9*) displayed a significant negative homozygous effect (-13.80%).

**Table 2.**
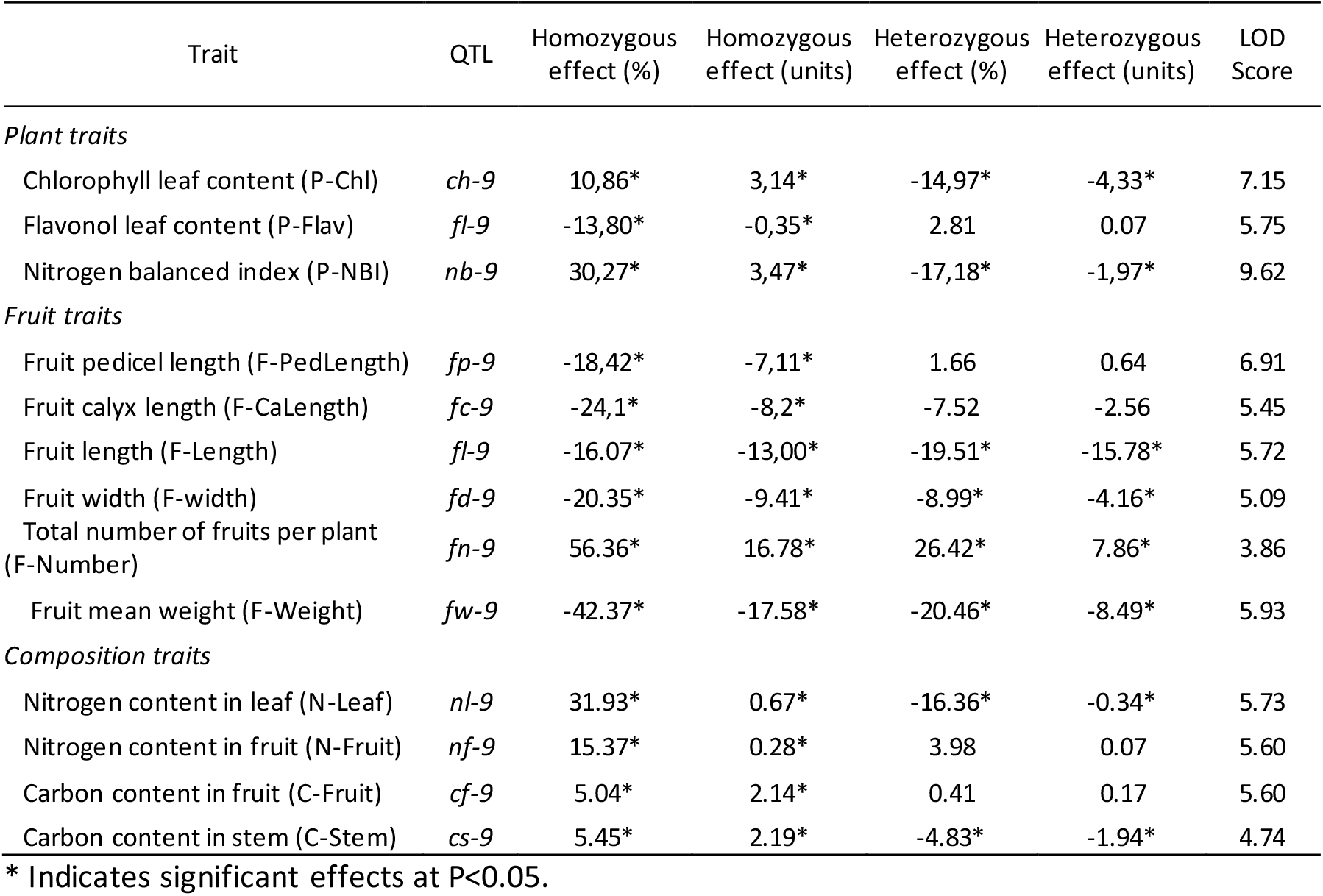
Putative QTLs detected corresponding to an introgression between 6.43 and 27.90 Mb of chromosome 9 of *S. insanum* with QTL name, homozygous and heterozygous effects, and LOD scores.

Regarding QTLs of fruit traits, a significant decrease in fruit pedicel and calyx length (F-PedLength and F-CaLength) was observed due to the effect of the homozygous allele from the wild species on QTLs *fp-9* and *fc-*9, respectively. Additionally, QTLs associated with fruit length (*fl-9*), width (*fd-9*) and fruit mean weight (*fw-9*), exhibited a significant negative effect for both homozygous and heterozygous alleles (Table2). The total number of fruits per plant-related QTL (*fn-9*) showed a significantly positive effect for both homozygous (56.36%) and heterozygous (26.42%) alleles, compared to the allele from the cultivated eggplant.

Nitrogen content in leaves and fruits (N-Leaf and N-Fruit), along with carbon content in fruits and stem (C-Fruit and C-Stem), were associated with QTLs *nl-9, nf-9, cf-9*, and *cs-9* respectively, demonstrating a significant positive effect of homozygous allele from the wild species (Table 2). However, QTLs *nl-9* and *cs-9* showed a negative significant heterozygous effect (-16.36% and -4.83%, respectively).

## 4. Discussion

The incorporation of CWR introgressions into populations with a cultivated background, such as ABs and ILs, enables the development of crop varieties with improved adaptive properties against abiotic stresses related to climate change (Prohens et al., 2017). Previous studies have made efforts to enhance understanding of N use in eggplant, aiming at developing new cultivars with improved N efficiency (Mauceri et al., 2020; Rosa-Martínez et al., 2023; Villanueva et al., 2021, 2023a). In this study, introgression lines derived from two eggplant CWRs, *S. dasyphyllum* and *S. insanum*, containing genomic regions potentially interesting for adaptation under low N conditions were evaluated together with segregating hybrids derived from them.

In a previous study, three sets of ABs of eggplant were evaluated under the same low N conditions (Villanueva et al., 2023a). The findings of that study revealed the identification of five QTLs located on a genomic region of chromosome 2 of ABs of *S. dasyphyllum* and an additional five QTLs on a genomic region of chromosome 9 of ABs of *S. insanum*. Traits associated with QTLs on chromosome 2 for ABs of *S. dasyphyllum* included chlorophyll leaf content, aerial plant biomass, yield, fruit pedicel length and fruit mean weight. For the QTLs located on chromosome 9 in *S. insanum* ABs, the associated traits include flavonol leaf content, nitrogen balanced index, fruit mean weight and nitrogen content in leaves and stems.

In our study, lines with introgressions at the same location on chromosomes 2 and 9 from *S. dasyphyllum* and *S. insanum* (IL-M1-D2 and IL-M5-I9), respectively, along with segregating hybrids derived from crosses between those lines and advanced backcrosses (ABs), cultivated under low N conditions allowed the evaluation and validation of previously detected QTLs, examining the impact of genetic background and different allele dosage. Our results revealed significant differences between the evaluated line sets, except for traits related to plant vigor and NUE parameters. Notably, both segregating hybrids resulting from two crosses (MEL1 X IL^Het^-M5-I9 and IL-M1-D2 X IL^Het^-M5-I9) exhibited no significant differences for any of the traits evaluated. This finding reveals that in our study the introgression from *S. dasyphyllum* on chromosome 2 did not manifest a discernible impact on the phenotypes of the segregating hybrids, revealing the importance of yearly differences in the expression of this QTL. However, within these segregating hybrids, a wider distribution ranges for all traits were observed, highlighting the potential of these materials to enhance variability in eggplant and assess the potential effects of genetic background. As occurred in other eggplant hybrids (Rehman et al., 2021), negative heterosis for specific traits was observed in segregating hybrids derived from crosses between the genetic backgrounds of *S. melongena* MEL1 and MEL5. The differences observed among materials suggest an increase in variation with respect to the MEL1 and MEL parents, emphasizing their potential significance for breeding.

The PCA and PLS analyses displayed a distinctive distribution pattern within the evaluated materials. Notably, a distinct grouping of individuals emerged from the IL of *S. insanum* (IL-M5-I9), differentiating it from other materials due to their association with traits related to high chlorophyll and nitrogen content. Conversely, individuals from the IL of *S. dasyphyllum* (IL-M1-D2) exhibited very slight differences with their corresponding parental line, *S. melongena* MEL1. In addition, both segregating hybrids were positioned close to each other. As stated before, these findings suggest a limited effect of the introgression of *S. dasyphyllum* on chromosome 2, with no significant differences detected.

The identification of 13 putative QTLs on a genomic region of chromosome 9 of *S. insanum* demonstrates the significant impact of this region on diverse traits. This finding allowed the validation of a set of QTLs for traits such as flavonol content in leaves, nitrogen balanced index, mean fruit weight, and leaf nitrogen content, as previously identified in Villanueva et al. (2023a) in segregating ABs. A new QTL was also identified for chlorophyll leaf content, a trait closely related to others (Cerovic et al., 2012). Also, QTLs associated with fruit size and morphology on chromosome 9 have been consistently found in various eggplant populations in different studies (Doganlar et al., 2002; Frary et al., 2014; Rosa-Martínez et al., 2023; Wei et al., 2020). Another novel QTL was identified for the total number of fruits per plant in this region. Finally, novel QTLs were detected in chromosome 9 for nitrogen content in the fruit, as the only one detected so far was in introgression lines of *S. incanum* on chromosome 12 (Rosa-Martínez et al., 2023), and carbon content in fruit and stem. Our results provide further evidence that environmental conditions, their interaction with genotypes, and genetic background can profoundly influence the phenotypic expression and stability of eggplant QTLs (Mangino et al., 2020; Mistry et al., 2016). In this way, the validation of detected QTLs in multiple locations and populations becomes crucial, considering the importance of the conditions established in the study (Diouf et al., 2018; Frary et al., 2014).

## 5. Conclusions

The evaluation of introgression and segregating hybrids revealed significant phenotypic variations in the materials evaluated, suggesting the potential of these materials for eggplant breeding for more sustainable agriculture practices. Notably, we confirmed that a genomic region from chromosome 9 of *S. insanum* may have a large impact on the adaptation of eggplant to low N conditions, as it contains a large number of QTLs, some of which are novel and that can be of interest for eggplant breeding. In contrast, we could not validate a genomic region from chromosome 2 of *S. dasyphyllum* that had been associated with chlorophyll content in leaves, aerial plant biomass and yield. The findings highlight the importance of validating eggplant QTLs across diverse conditions and populations, employing techniques like evaluating the genomic regions under different backgrounds, and considering the complex interactions influencing quantitative trait expression, including environmental factors, genotypes, and genetic backgrounds.

## Data availability

Relevant data can be found within the paper. All data of this study are available from the corresponding author upon request.

## Acknowledgements

This work was supported by the project SOLNUE in the framework of the H2020 call SusCrop - ERA-Net (ID#47) and funded by Agencia Estatal de Investigación (PCI2019-103375), by the Spanish Ministerio de Ciencia e Innovación, Agencia Estatal de Investigación, ( grants RTI2018-094592-B-I00 from MCIU/AEI/ FEDER, UE, and PID2021-128148OB-I00, funded by MCIN/AEI/10.13039/501100011033/ and “ESF Investing in your future”), and by Conselleria d’Innovació, Universitats, Ciència i Societat Digital of the Generalitat Valenciana (grant CIPROM/2021/020). The Spanish Ministerio de Ciencia e Innovación, Agencia Estatal de Investigación, and Fondo Social Europeo funded a predoctoral fellowship to Gloria Villanueva (PRE2019-089256). Pietro Gramazio is grateful to Spanish Ministerio de Ciencia e Innovación for a post-doctoral grant (RYC2021–031999-I) funded by (MCIN/AEI /10.13039/ 501100011033) and the European Union through NextGenerationEU/ PRTR.

